# Identification of mRNAs that undergo stop codon readthrough in *Arabidopsis thaliana*

**DOI:** 10.1101/2021.11.09.467898

**Authors:** Sarthak Sahoo, Divyoj Singh, Anumeha Singh, Sandeep M Eswarappa

## Abstract

A stop codon ensures termination of translation at a specific position on an mRNA. Sometimes, termination fails as translation machinery recognizes a stop codon as a sense codon. This leads to stop codon readthrough (SCR) resulting in the continuation of translation beyond the stop codon, generating protein isoforms with C-terminal extension. SCR has been observed in viruses, fungi, and multicellular organisms including mammals. However, SCR is largely unexplored in plants. In this study, we have analyzed ribosome profiling datasets to identify mRNAs that undergo SCR in *Arabidopsis thaliana*. Analyses of the ribosome density, ribosome coverage and three-nucleotide periodicity of the ribosome profiling reads, in the mRNA region downstream of the stop codon, provided strong evidence for SCR in mRNAs of 144 genes. This process generates putative peroxisomal targeting signal, nuclear localization signal, prenylation signal, transmembrane helix and intrinsically disordered regions in the C-terminal extension of several of these proteins. Gene ontology (GO) functional enrichment analysis revealed that these 144 genes belong to three major functional groups - translation, photosynthesis and abiotic stress tolerance. Finally, using a luminescence-based assay, we experimentally demonstrate SCR in representative mRNAs belonging to these functional classes. Based on these observations, we propose that SCR plays an important role in plant physiology by regulating the protein localization and function.

**AUTHOR SUMMARY:** Protein synthesis executed by macromolecular complexes, termed ribosomes, starts and stops at specific locations on a messenger RNA (mRNA). This fidelity is critical for the normal functioning of cells. However, sometimes ribosomes don’t stop translation at the stop signal (termed stop codon) on an mRNA resulting in longer proteins with properties different from those of the canonical shorter protein. This process called stop codon readthrough (SCR) has been observed in viruses, fungi, and multicellular organisms including mammals. However, it remains largely unexplored in plants. In this study, we report evidence of SCR in 144 genes of *Arabidopsis thaliana*, a small flowering weed widely used as a model system to study plant biology. These genes are involved in protein synthesis, photosynthesis and stress tolerance in plants. We have also experimentally demonstrated SCR in a few genes that represent these functional classes. Our analysis shows that SCR can change the localization and functional properties of these proteins. We propose that SCR plays an important role in plant physiology.

## INTRODUCTION

A stop codon (UGA/UAA/UAG) on an mRNA signals the translating ribosomes to terminate the process of translation. However, in certain mRNAs, ribosomes fail to terminate at the canonical stop codon and continue translation till the next in-frame stop codon. This is caused by recoding of stop codons by a near-cognate tRNA or a suppressor tRNA. This process of stop codon readthrough (SCR) generates protein isoforms with extended C-terminus, thus contributing to proteome expansion [1]. Because of the extended C-terminus, the protein isoform generated by SCR can be different from the canonical isoform in terms of its localization, function, or stability [2–5]. Since this process occurs at the translational level, SCR enables cells to swiftly respond to environmental cues.

SCR has been observed in bacteria, yeast, insects, mammals, and viruses. It is well-studied in plant viruses [6,7]. For example, tobacco necrosis virus-D (TNV-D) expresses its polymerase, and potato leafroll virus generates a minor capsid protein by SCR [8,9]. It enables viruses to maximize the coding potential of their compact genome. Since plant viruses utilize the translation machinery of the host, it is likely that some plant mRNAs also undergo SCR. However, so far, there is only one report of a plant mRNA undergoing SCR. *Arabidopsis* eRF1-1 mRNA undergoes SCR, which regulates its expression by protecting the mRNA from non-sense mediated decay [10]. A wide range of other translation regulation mechanisms have been observed during plant development, light-dark cycle, viral infections, and environmental stresses [11]. A genome-wide analysis of SCR, which is also a translation regulation mechanism, is lacking to understand its role in plant physiology.

Ribosome profiling technique, based on the deep sequencing of ribosome protected RNA fragments, has revolutionized our understanding of the process of translation and its regulation. Because it reveals ribosome-occupied regions on an mRNA, ribosome profiling has the potential to identify novel translation events such as SCR [12]. In this study, we analyzed ribosome profiling datasets and identified 144 genes of *Arabidopsis thaliana* as targets of SCR. Further, we experimentally confirmed this process in 4 candidate genes.

## RESULTS AND DISCUSSION

### Selection and curation of ribosome profiling datasets

The presence of translating ribosomes after the canonical stop codon of an mRNA strongly indicates SCR [12]. We analyzed ribosome profiling data generated using *A. thaliana,* which are available at Sequence Read Archive (SRA) of National Center for Biotechnology Information (NCBI). We retrieved 14 *A. thaliana* ribosome profiling datasets from SRA and processed them as described in Methods.

Ribosomal footprints obtained from translating ribosomes exhibit frame bias. i.e., they show a fixed distribution of reads across the three frames of the coding sequence. This three-nucleotide periodicity (or phasing) is a sign of translation on the corresponding region of an mRNA (in our case, the proximal 3′UTR). This kind of spatial resolution along mRNAs is required in ribosome profiling datasets to claim unusual translation events such as SCR. Therefore, we first analyzed the three-nucleotide periodicity profile of the ribosome profiling datasets. We chose 9 ribosome profiling datasets based on a clear three-nucleotide periodicity of ribosome profiling (ribo-seq) reads corresponding to the coding sequences of all genes. A representative profile is shown in Fig 1A.

**Figure 1.**
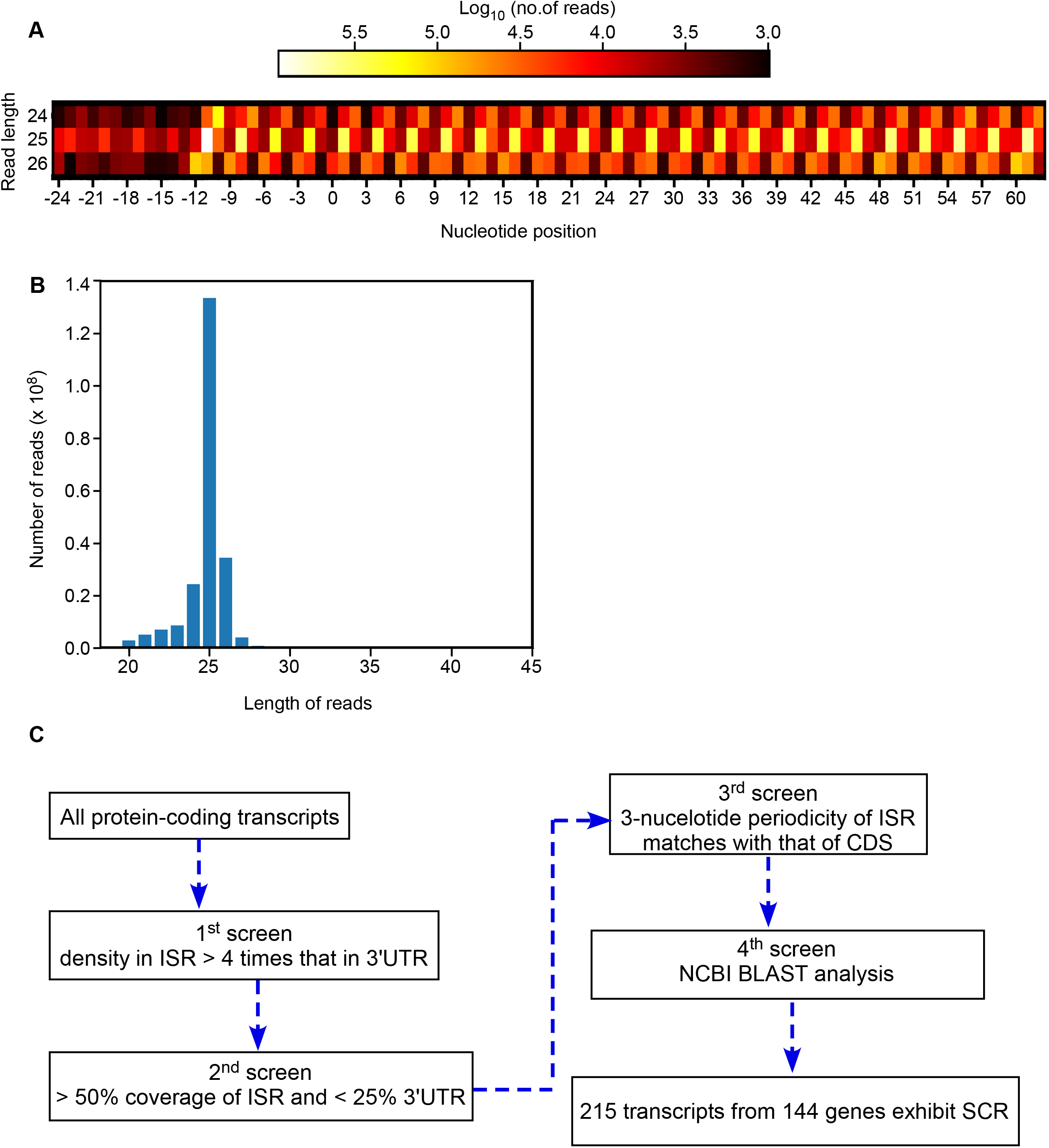
Selection and analysis of ribosome profiling datasets. (A) Heat map showing the three-nucleotide periodicity profile of the dataset SRP074840. Ribosomal footprints on all coding sequences were analyzed to get this profile for read lengths 24, 25, and 26. Reads were assigned to three frames based on which frame the first nucleotide aligns with on an mRNA sequence. The start codon ATG is indicated by the position 0 on the x-axis. (B) Distribution of ribo-seq read lengths. The graph shown is from the dataset SRP074840 (C) Flow chart showing the four-level screening method to identify mRNAs that show SCR in *A. thaliana*. Another flow chart with more details is shown in Fig S2.

These 9 datasets were derived from seedlings, root, shoot, flower, and from a cell line of *A. thaliana* (Table S1) [13–20]. We then analyzed the length distribution of the reads in each dataset. Length distribution was consistent with footprints of 80S ribosomes, which is a signature of translation. A representative length distribution is shown in Fig 1B. Based on this distribution, we chose the reads of 3 most abundant lengths for further analyses.

After removing the reads that map onto non-coding RNAs, we aligned the rest of the reads with *A. thaliana* protein-coding mRNAs. Only those reads that align 100% (i.e., without any mismatch) with an mRNA were considered for the analysis. As our aim was to identify SCR, we focused on the ribosome footprints in the proximal part of the 3′UTR - from the canonical stop codon to the downstream in-frame stop codon (Fig S1). This region was termed inter-stop codon region (ISR). mRNAs without downstream in-frame stop codon were not included in the analysis.

### 215 mRNAs from 144 genes of *A. thaliana* show evidence of SCR

We subjected the mRNAs of *A. thaliana* to a stringent four-level screening to identify the targets of SCR (Fig 1C and Fig S2). It is important to distinguish ribosome profiling (ribo-seq) reads due to SCR from reads resulting from non-translating events [21]. This was achieved by comparing the ribosome densities in different regions of an mRNA. The average ribosome density in the 3′UTR (untranslated region) of mRNAs indicates ribosome occupancy due to events not related to translation. mRNAs that showed at least 4-fold higher ribosome density in the ISR compared to the rest of the 3′UTR were considered for further analysis. 1144 mRNAs were identified in this first level of screening (Fig S2 and Table S1).

It is possible that the increased ribosome density can be due to a specific segment of the ISR with a strong RNA structure or a strong interaction with a protein (or any *trans*-acting molecule). To exclude such events, we subjected the mRNAs to a coverage-based filtering. We eliminated mRNAs with < 50% coverage in the ISR and > 25% coverage in the rest of the 3′UTR. 550 mRNAs satisfied this criterion and all of them had at least one ribo-seq read spanning the canonical stop codon (Fig S2 and Table S1).

The three-nucleotide periodicity profile of the ribo-seq reads assigned to ISR, similar to that of the reads assigned to the coding sequence (CDS), provides a strong evidence for SCR. 236 mRNAs satisfied this third level of screening. It is possible that the remaining mRNAs may include ribosomal frameshifting (also known as translational frameshifting) candidates. We did not pursue frameshifting events as the focus of our study was SCR, where the frame of the ribosomes translating the ISR is same as that in the CDS. The ribosome density profile and the three-nucleotide periodicity profile of the ISRs of four representative genes - *RPS15AD, CURT1B, CAM1* and *MUB6* - are shown in Fig 2.

**Figure 2.**
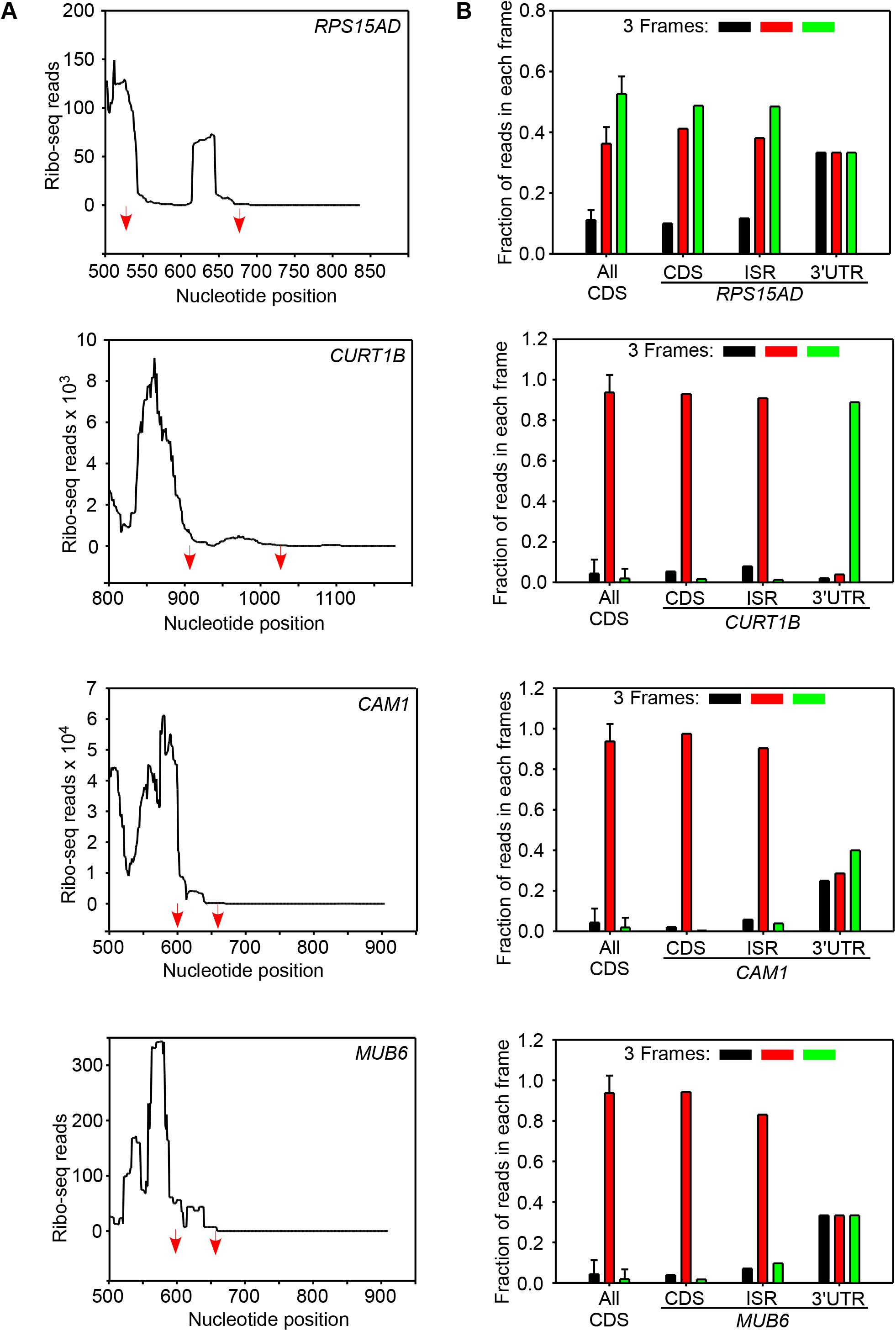
Ribosomal density and three-nucleotide periodicity at the ISR of 4 SCR-positive mRNAs –*RPS15AD*, *CURT1B*, *CAM1*, *MUB6*. (A) Graphs showing ribo-seq reads in the ISR of four genes. Red arrows indicate the position of the two stop codons. Some parts of the coding sequence and the 3′UTR are also shown for comparison. (B) Three-nucleotide periodicity. Graphs show fraction of ribo-seq reads in three translation frames. The three-nucleotide periodicity profile of coding sequences of all protein-coding genes is shown for comparison (All CDS). The data shown in (A) and (B) are from the dataset SRP074840. CDS, coding sequence; ISR, inter-stop codon region; UTR, untranslated region.

The mRNA sequences of *A. thaliana* were retrieved from Ensembl Plants. The annotation of the coding sequence on mRNAs can vary across the databases, providing false-positive evidence for SCR. To rule this out, we performed BLAST analysis for the peptides encoded by the ISRs of 236 mRNAs that passed the screening described above, against NCBI’s protein database for *A. thaliana*. We found 21 matches, which were eliminated in this fourth level screening.

Thus, 215 mRNAs encoded by 144 genes of *A. thaliana* passed our stringent four-level screening, and they were designated as SCR-positive genes (Fig S2 and Table S2). The average ribosome density in the ISR increased with each screening step (Fig 3A). Also, the average ribosome density in the ISR of SCR-positive mRNAs was 10-fold higher compared to the same in all mRNAs. This difference was not observed in the ribosome density of the 3′UTR excluding the ISR (Fig 3B).

**Figure 3.**
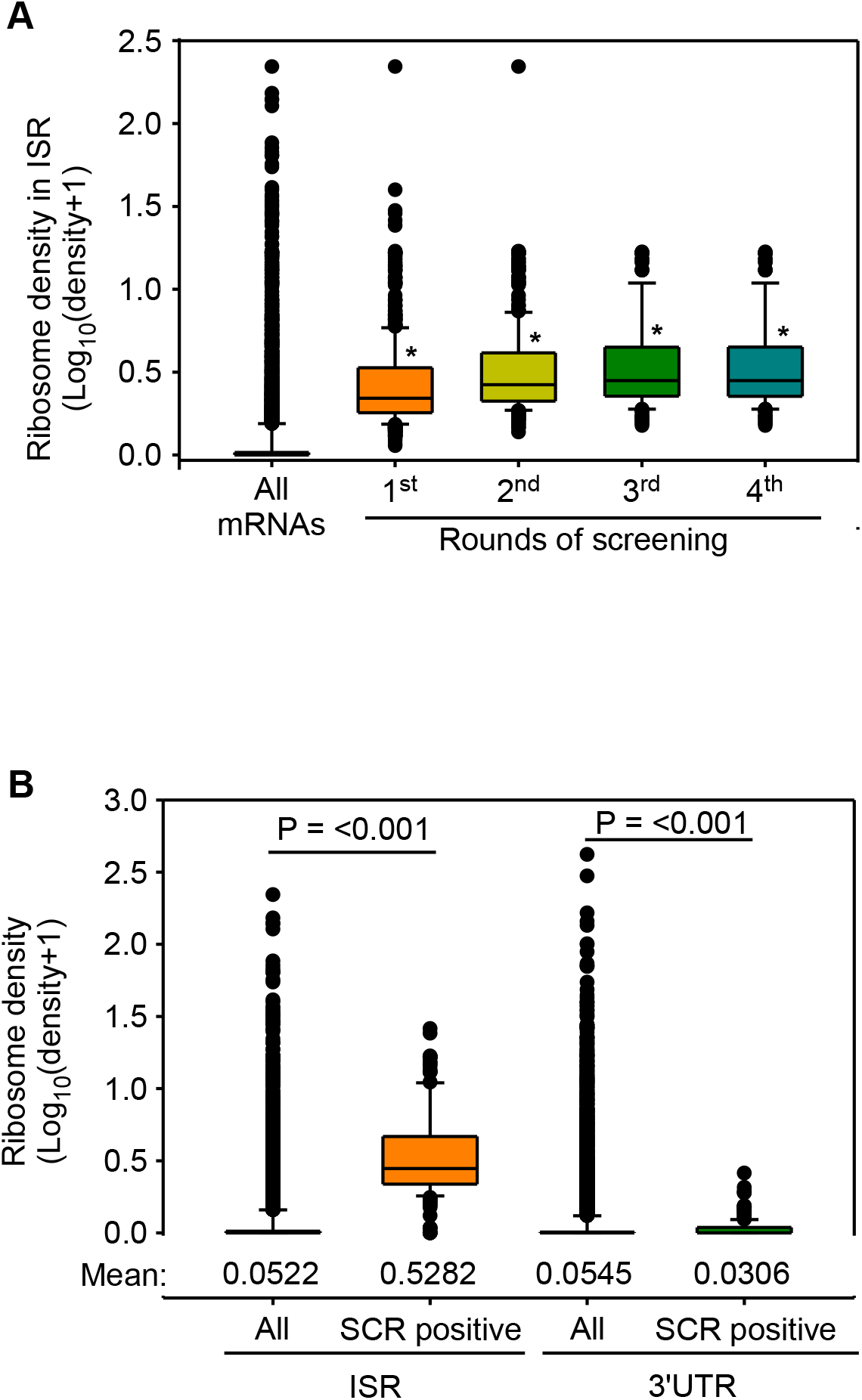
Ribosomal density in the ISR of SCR-positive mRNAs. (A) The graph shows the increase in ribosome density in the ISRs after each round of screening. *, P < 0.001 Mann-Whitney Rank Sum Test (compared to ‘All mRNAs’). (B) Graph shows the comparison of ribosomal density in the ISR vs that in the 3′UTR. This comparison is shown for SCR-positive mRNAs and for all mRNAs. P values were calculated using Mann-Whitney Rank Sum Test. Numbers at the bottom of the graph indicate the mean value. The box represents 25% and 75% values, and the horizontal line within the box shows the median value. The analysis shown is for the dataset SRP074840.

These results show that our screening methods were able to identify translational event immediately after the stop codon, which constitutes SCR. Though translational frameshifting could also result in increased ribosomal density in the 3′UTR, the three-nucleotide periodicity-based 3rd screen will remove frameshifting events as described above. We have not allowed a single mismatch while assigning reads to different regions of an mRNA. Also, all SCR-positive candidates have at least one ribosome profiling read mapping on to the region spanning the canonical stop codon (the junction of the coding sequence and the ISR). These two conditions rule out RNA editing and polymorphism at the stop codon as reasons for ribosome footprints after the stop codon.

In our analyses, we have excluded genes which have ISRs and/or 3′UTRs < 45 nucleotides and genes with < 30 reads in their ISR. Also, our method does not reveal candidates that undergo SCR under specific physiological or pathological conditions not included in the ribosome profiling studies. Hence, 144 SCR-positive genes is an underestimate; it is likely that more mRNAs undergo SCR in *A. thaliana*. For example, we did not observe any ribo-seq footprints in the ISR of *eRF1-1*, which has been demonstrated to undergo SCR in *A. thaliana* [10].

### The stop codon TGA is enriched in SCR-positive genes

Since the identity of the stop codons can influence the efficiency of translation termination [22], we examined the distribution of the three stop codons among the SCR-positive mRNAs at their canonical termination position. We observed a 25% higher occurrence of TGA stop codon in SCR-positive mRNAs compared to the expected frequency (Fig 4A). Interestingly, TGA is the leakiest among the three stop codons, which facilitates the process of SCR.

**Figure 4.**
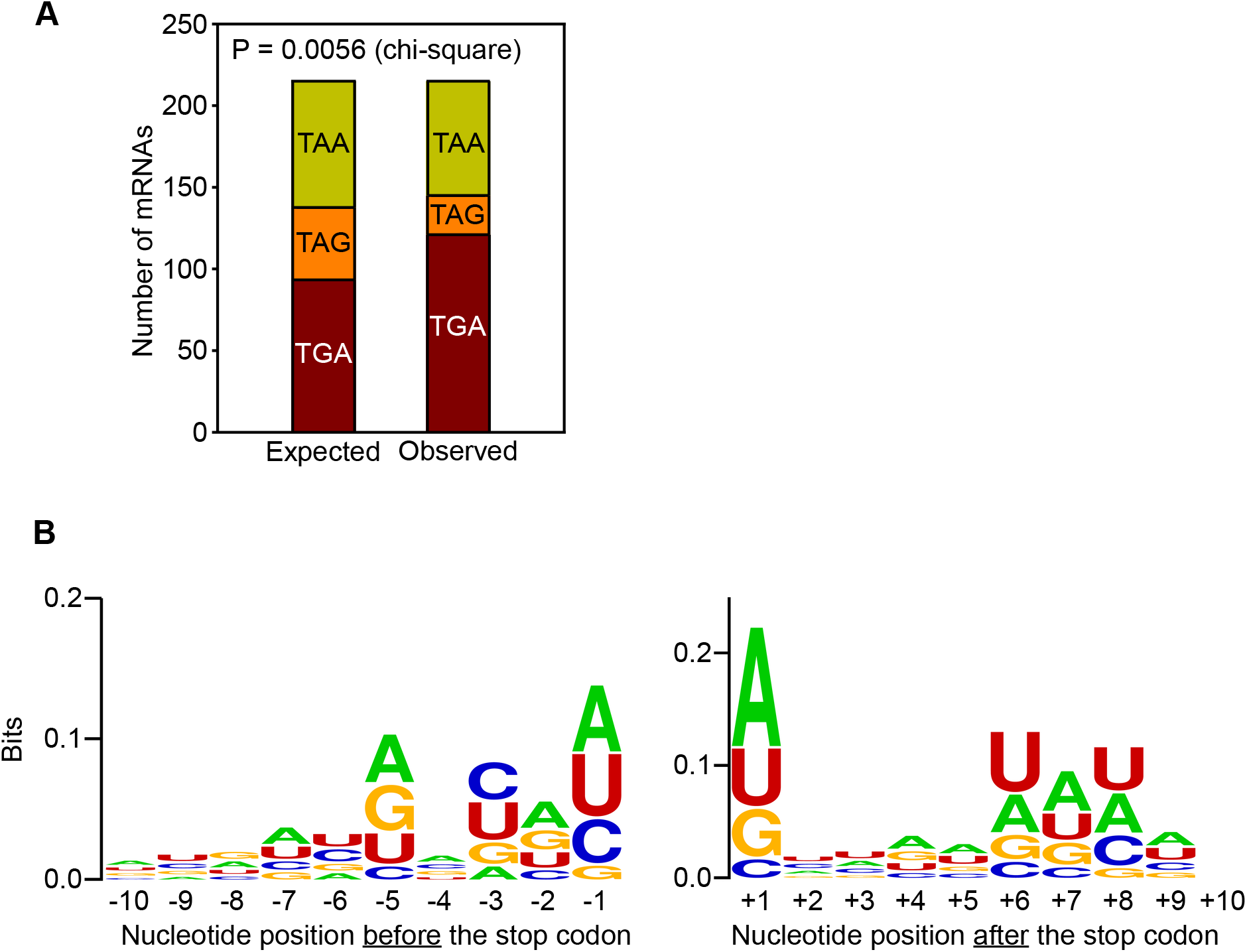
The canonical stop codon and its context in SCR-positive mRNAs. (A) Distribution of the three stop codons in SCR-positive mRNAs. Expected values were obtained based on their occurrence in all mRNAs of *A*. *thaliana*. (B) Sequence logo of the stop codon context of SCR-positive mRNAs. The analysis was performed using WebLogo.

**Figure 5.**
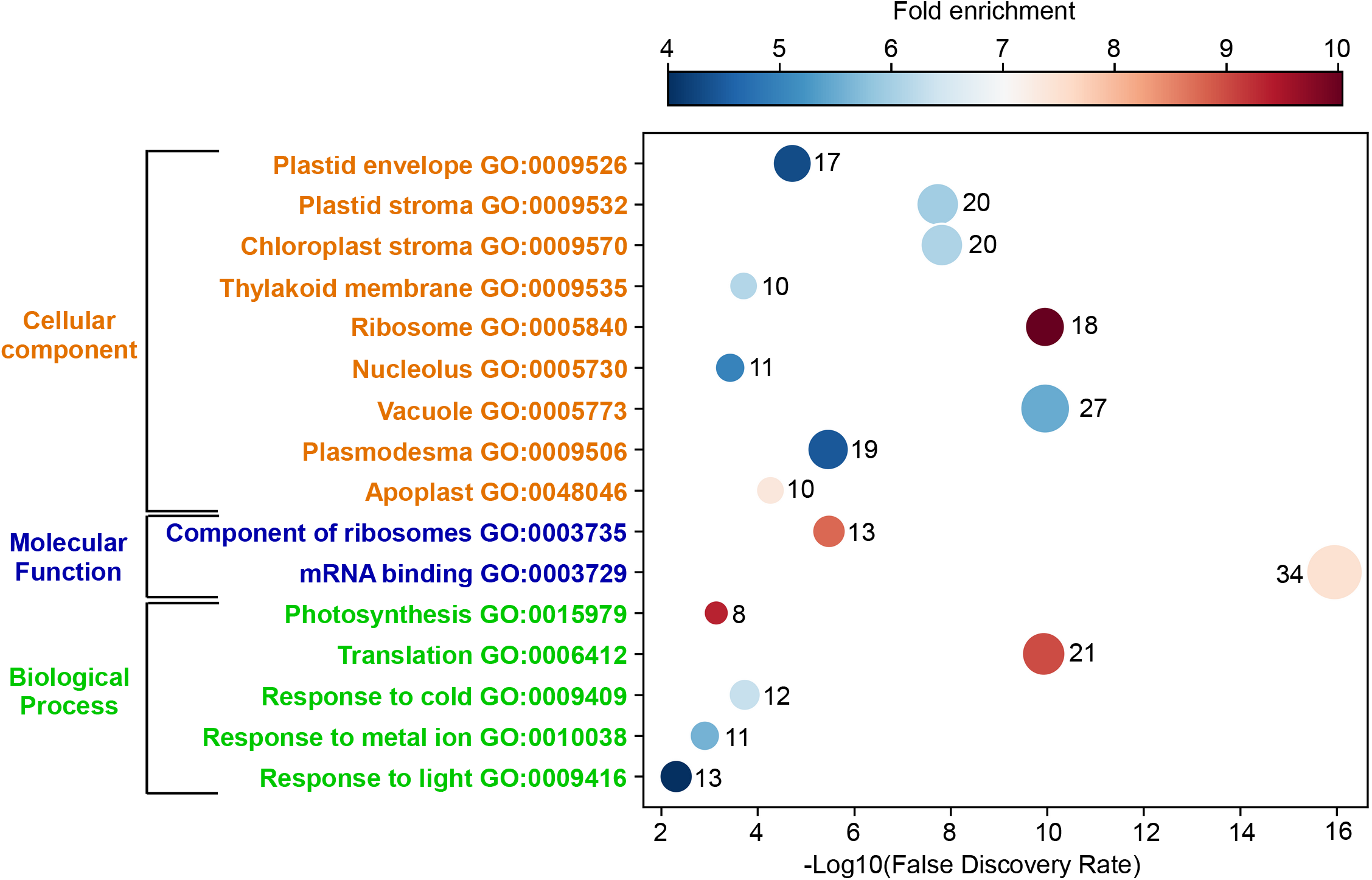
Gene ontology analysis of SCR-positive genes of *A. thaliana*. Results of gene ontology (GO) functional enrichment analysis on SCR-positive genes using PANTHER web server. The X-axis shows false discovery rate and the Y-axis shows multiple functional classes enriched in SCR-positive genes. Color of the circle indicates fold enrichment. The number of SCR-positive genes showing enrichment in a functional group is shown next to the circle. Size of the circle is proportional to this number. SCR-positive genes showing more than 4-fold enrichment are shown here.

The context of stop codons, especially the nucleotides immediately before (−1) and after (+1) the stop codon, can also influence the efficiency of translation termination [23]. Hence, we examined if there are any conserved sequences around the stop codon in SCR-positive mRNAs. We used WebLogo, a sequence logo generator, to visualize the extent of conservation around the stop codon [24]. Here, the height of the stack indicates the extent of conservation at that particular position. Interestingly, nucleotides just before (−1) and after (+1) and the stop codon showed higher conservation compared to other positions. A and U were more frequently observed in these positions than the other two nucleotides (Fig 4B). These conserved residues are possibly important to provide SCR-permissive context in SCR-positive mRNAs.

### Gene ontology analysis: Genes involved in translation, photosynthesis, and stress response are enriched in SCR-positive genes

To gain some insight into the functional significance of SCR in *A. thaliana*, we performed gene ontology (GO) functional enrichment analysis on 144 SCR-positive genes using PANTHER web server [25]. Among biological processes, we observed that genes involved in translation, photosynthesis, and abiotic stress response were enriched in the list of SCR-positive genes. For instance, 21 genes involved in translation were part of this list. In consistence with this, genes encoding proteins localized in ribosomes and nucleolus (site of ribosome assembly) were enriched. With respect to molecular functions, there was an enrichment of genes encoding components of ribosomes and mRNA-binding proteins. Interestingly, 34 SCR-positive genes encode RNA-binding proteins. Proteins encoded by 20 of them localize in chloroplast and 18 of them are ribosomal proteins. Together, these observations indicate that SCR could play an important role in regulating the process of translation, photosynthesis, and abiotic stress response in *A. thaliana*.

### SCR can change the localization of the proteins

SCR has been shown to change the localization of the protein product in some cases. For example, the SCR product of mammalian *MTCH2* is localized to the cytoplasm while the canonical isoform is a mitochondrial membrane protein [3]. SCR products of mammalian malate dehydrogenase and lactate dehydrogenase have a peroxisomal targeting sequence (PTS) at the C-terminus, which directs them to peroxisomes. However, the canonical isoforms are found in the cytoplasm or mitochondria [5,26,27]. Since PTS is usually found at the C-terminus of a protein, SCR provides a mechanism to generate peroxisomal protein isoforms. We analyzed the extended C-termini of 144 *A. thaliana* proteins potentially generated after SCR for a possible PTS using PredPlantPTS1 tool [28]. We found four of them with a PTS at their extended C-terminus - AT1G09310 (unknown function), *CHS* (encodes chalcone synthase), *AGP15* (encodes an arabinogalactan protein) and *GGH2* (encodes a gamma-glutamyl hydrolase). None of the canonical isoforms of these four gene-products is known to be located in the peroxisomes (Table 1). Thus, SCR can drive the protein isoforms generated by these four genes to peroxisomes and regulate their functions.

**Table 1.** List of SCR-positive genes whose products exhibit peroxisomal targeting sequence after SCR^$^.

| Gene | Function* | Location of the canonical protein* | Peptide encoded by the ISR<sup>#</sup> |
| --- | --- | --- | --- |
| AT1G09310 | Unknown | apoplast, cytosol, extracellular region, nucleus | SAQLQQIKETRIFKCT**SRI** |
| AT5G13930<br>(*CHS*) | chalcone synthase involved in the biosynthesis of flavonoids | cytoplasm, endoplasmic reticulum, nucleus, plant-type vacuole membrane | ERLPSICLPTY**AKL** |
| AT5G11740<br>(*AGP15*) | arabinogalactan protein | plasma membrane | VTVMVISYRDCFCGIGHSS<br>LFVVSCVFR**SSL** |
| AT1G78680<br>(*GGH2*) | a gamma-glutamyl hydrolase acting specifically on monoglutamates. | Vacuole | NGGFCRIGYDEVYIFTQQR**SLL** |

<sup>\$</sup> The analysis was done using the PredPlantPTS1 tool
\*Function and location information were obtained from The Arabidopsis Information Resource (TAIR)
<sup>#</sup> Experimentally verified plant PTS (peroxisome targeting sequence) tripeptides are highlighted in red

Nuclear localization signal (NLS) at the C-terminus of the SCR products can drive them to the nucleus. We searched for the presence of NLS in the ISR of the 144 SCR products using SeqNLS and NLStradamus [29,30]. Three of them showed strong NLS with a score > 0.7 - *CURT1B, KCS12* and *AT5G56200*. Among these, the canonical protein isoforms of *CURT1B* (The P subunit of Photosystem I) and *KCS12* (3-ketoacyl-CoA synthase) are not localized in the nucleus. Our analysis predicts that their SCR isoforms are localized in the nucleus, possibly with a moonlighting function (Table 2).

**Table 2.** List of SCR-positive genes whose products exhibit Nuclear localization signal (NLS) in the extended C-terminus^$^.

| Gene | Function* | Peptide encoded by ISR<sup>#</sup> |
| --- | --- | --- |
| AT2G46820<br>(*CURT1B*) | The P subunit of Photosystem I | IKGGRRRRRAFLRPFMNWNE<br>GYQKNLTQRPRPSFNLSFL<br>(0.727) |
| AT2G28630<br>(*KCS12*) | 3-ketoacyl-CoA synthase<br>(involved in the biosynthesis of<br>very long chain fatty acids) | NVYAQKRKRKRKNNT<br>RIELVKTCLAIGKPNKCV<br>(0.895) |
| AT5G56200 | Encodes a transcription factor<br>expressed in the female<br>gametophyte. | ETYICKQVIFLTLKKKKTKK<br>(0.862) |

<sup>\$</sup>The analysis was done using SeqNLS and NLStradamus
\*Function information was obtained from The Arabidopsis Information Resource (TAIR)
<sup>#</sup>NLSs are highlighted in red and the numbers indicate the SeqNLS score

We then searched for a transmembrane helix at the C-terminus of 144 SCR isoforms using TMHMM Server v. 2.0 [31], as this can be a mechanism to change the localization of the protein to a membrane. 23 of the SCR products showed a transmembrane helix at their extended C-terminus (Table S3). The canonical isoforms of 17 of these 23 genes are not known to be membrane proteins. Our analysis suggests that SCR can potentially regulate their function by driving them to the cell membrane. We also searched for endoplasmic reticulum retention signal, KDEL, in the peptides encoded by the ISR of SCR-positive genes. None of them possess this signal.

Isoprenylation is a post-translational modification that occurs at a cysteine residue in the C-terminus. This modification can lead to anchoring of the protein to the cell membrane. We analyzed the peptides encoded by the ISR of 144 SCR-positive genes for potential prenylation signal using the PrePS tool [32]. The peptide encoded by the ISR of metallothionein 1C (MT1C; AT1G07610) showed a potential prenylation signal at its C-terminus – RNYQHGLKPRKMGKK<u>CVLC</u>. CVLC is the predicted prenylation signal. Metallothioneins are cysteine-rich proteins required for tolerance to heavy metals, an abiotic stress. Prenylation of the extended isoform of MT1C can potentially alter its localization from the cytosol to membranes regulating its function.

In case of mammalian *VEGFA* and *AGO1*, SCR results in a C-terminus with intrinsically disordered region (IDR). This changes the functional properties of their SCR isoforms [2,4]. We analyzed the products of 144 SCR-positive genes for possible IDRs at the C-terminus using the IUPred2A tool [33]. We observed IDR in the C-terminus of the products of 6 SCR-positive genes. They are involved in the organization of cytoskeleton, redox homeostasis, fatty acid synthesis, auxin and hypersensitive response (Table 3). IDR in their C-terminus can potentially alter the functional properties of these six SCR products.

**Table 3.**
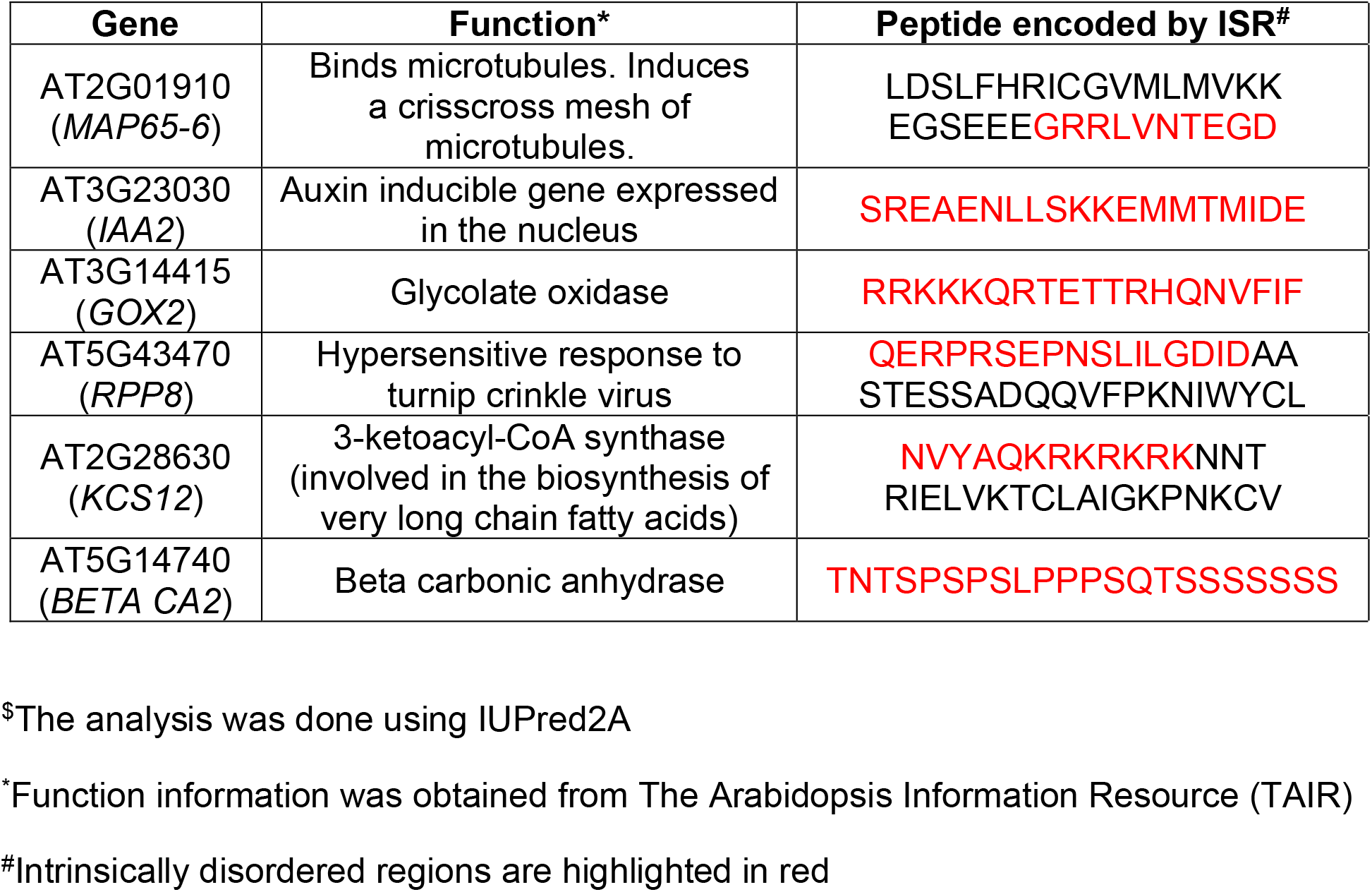
List of SCR-positive genes whose products exhibit intrinsically disordered regions in the extended C-terminus after SCR^$^.

### Experimental validation of SCR

We performed *in vitro* translation experiments using wheat germ extract to validate SCR in four genes: *RPS15AD* encodes a ribosomal protein; *CURT1B* encodes the P subunit of Photosystem I; *CAM1* encodes calmodulin, which is involved in abiotic stress response [34]. These three genes represent three functional classes that are enriched in SCR-positive genes - translation, photosynthesis, and abiotic stress response. We also selected one more gene, *MUB6* (encodes membrane-anchored ubiquitin-fold protein 6), which does not belong to any of these three classes, but is one of the 144 SCR-positive genes. As described above, ribosome footprints were observed after the stop codon in the ISR of these mRNAs. Also, the three-nucleotide periodicity of the ribosomal footprints on the ISR was comparable to that of the coding sequence, but not to that of the 3′UTR (Fig 3).

Luminescence-based SCR assays were performed as described previously [4]. We cloned the cDNAs of these genes upstream of and in-frame with the cDNA of firefly luciferase without its start codon. Luminescence will be observed only if the translation continues across the canonical stop codon of the test cDNA (Schematic in Fig 6). Thus, luminescence in these assays indicates SCR. *In vitro* transcription followed by *in vitro* translation using wheat germ extract revealed significant luminescence activity in mRNAs of all four genes, much above the background level. Constructs without the corresponding ISRs were used to know the background level of luciferase activity. A construct without a stop codon between the test cDNA and the firefly luciferase cDNA was used to measure the efficiency of SCR. This analysis revealed 8%, 50%, 25%, and 6.5% SCR in *RPS15AD*, *CURT1B, CAM1,* and *MUB6*, respectively.

**Figure 6.**
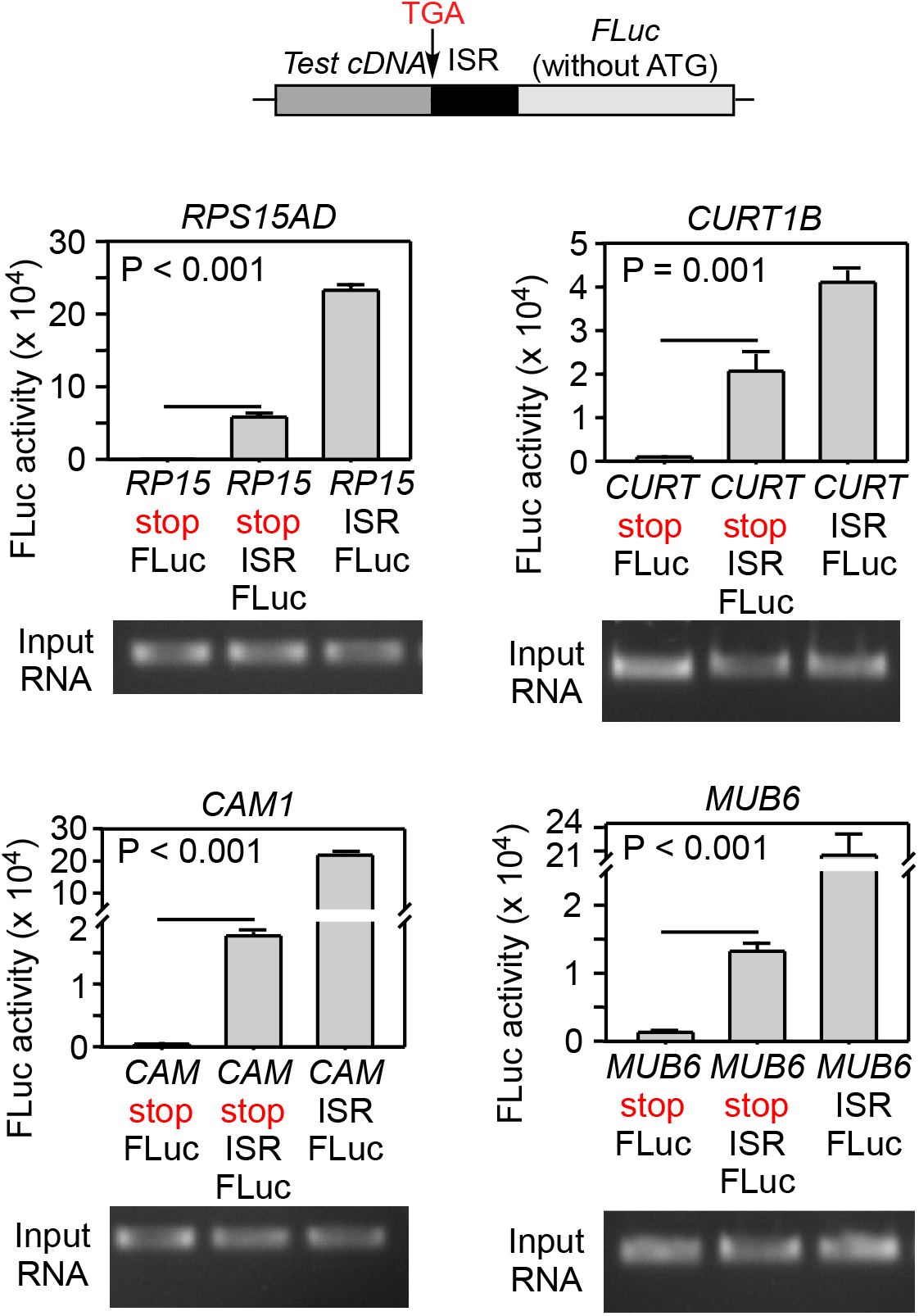
Experimental validation of SCR in four *A. thaliana* mRNAs – *RPS15AD*, *CURT1B*, *CAM1* and *MUB6*. Luminescence-based SCR assay. cDNA of a test gene along with the ISR was cloned upstream of and in-frame with the cDNA of firefly luciferase (FLuc) such that FLuc is expressed only if there is SCR across the stop codon of the test cDNA (see the schematic). Constructs were subjected to *in vitro* transcription followed by *in vitro* translation as described in Methods. Expression of Fluc was measured by its luminescence activity, which is shown in the graphs. Constructs without ISR were used to measure background signal (first bar), and constructs without any stop codon between the test cDNA and the FLuc were used to measure the maximum luminescence activity (third bar). Statistical significance (two-sided *P*-value) was obtained using Student’s t-test. Input RNA obtained by *in vitro* transcription is shown below the graphs.

Overall, our analysis of ribosome profiling datasets provides strong evidence for SCR in mRNAs of 144 genes of *A. thaliana*. Using a similar analysis of ribosome profiling data, mRNAs of 350 *Drosophila* genes and 42 human genes have been predicted to undergo SCR [12]. The advent of ribosome profiling technique has revealed previously unknown (or lesser-known) mechanisms of translational regulation, including SCR. Since this technique is based on experimentally generated ribosome footprints on mRNAs, it is superior to evolutionary conservation-based computational screening methods to detect SCR, which will miss SCR events that have emerged relatively recently during evolution. Furthermore, the nucleotide resolution of ribosome profiling enables us to decipher the frame of translation at the ISRs. The distribution of length of ribosome profiling reads will have a signature of 80S ribosome occupancy. These features are important to distinguish SCR from ribosomal frameshifting and non-translational events (e.g., protein binding and RNA structures) [35]. Thus, ribosome profiling is a powerful tool to identify SCR events at the transcriptome level.

It would be remarkable if SCR does change the properties of the proteins in multiple ways as predicted by our analyses – by introducing peroxisomal targeting signal, nuclear localization signal, prenylation signal, transmembrane helices and intrinsically disordered region in the ISR-encoded C-terminal extension. Other mechanisms such as post-translational modification and degradation, which cannot be predicted with high confidence, might also occur at ISR-encoded extensions. This is not very surprising as random peptide sequences have been shown to have functional motifs. For example, 1/5^th^ of randomly generated peptide sequences carry export signal in *Saccharomyces cerevisiae* [36]. Also, in another study involving *S. cerevisiae*, 8 out of 28 randomly generated peptides showed multiple organellar localization signals [37]. Therefore, for a gene the chances of acquiring novel functions by SCR are high.

As shown in mammalian and viral SCR processes, it is likely that the nucleotide sequence of ISR is responsible for driving the SCR via a *cis*-acting RNA motif or *trans*-acting molecule [2,4,38,39]. Thus, ISR likely possesses a dual function – driving the SCR and altering the properties of the SCR product. It will be interesting to study how natural selection will shape such genomic regions with constraints at both nucleotide (ability to induce SCR) and amino acid level (novel function).

The GO analysis suggests that SCR in *A. thaliana* influences three major physiological processes in plants – protein synthesis, photosynthesis and stress tolerance. Our *in vitro* translation experiments performed using a plant-based system show that the efficiency of SCR is much above the basal error rate, suggesting that these are programmed events with physiological consequences. This is consistent with various functional motifs identified in the C-terminal extensions. We anticipate that more studies will follow to characterize individual SCR events in order to understand the mechanism of SCR as well as its physiological significance in plants.

## Materials and Methods

### Curation of *A. thaliana* transcriptome

*A. thaliana* has 55,398 mRNAs derived from 27,655 protein-coding and 6,563 non-coding genes. (http://plants.ensembl.org/Arabidopsis_thaliana/Info/Annotation/#assembly). Sequences of the mRNAs were downloaded from Ensembl Plants. From this, we created a file containing sequences of rRNAs, tRNAs, snRNAs, snoRNAs, and miRNAs, which were later used to remove ribosomal footprints that aligned to these sequences. Using cDNA sequences (downloaded from the same source), we noted the positions of the start codon, the canonical stop codon and the first in-frame stop codon (if any). mRNAs with inter-stop codon region (ISR) < 45 nucleotides or rest of the 3′UTR < 45 nucleotides were removed. This is because sequences with shorter length will not give enough statistical power to draw any conclusions from the ribosomal density differences between them (i.e., ISR and 3′UTR). mRNAs whose ISR sequence was matching with > 24 nucleotide sequence of any other coding sequence were removed. This was done because reads cannot be mapped onto an ISR if its sequence matches with a coding sequence. After these filtrations, we were left with 14,732 protein coding mRNAs for our analysis.

### Preprocessing and sequence alignment of ribosome profiling datasets

Sequence Read Archive (SRA)-formatted ribosome profiling datasets of 9 studies on *A. thaliana* were downloaded from SRA (Table S1). They were converted to FASTQ format files using the prefetch and fastq-dump command of SRAToolkit (https://github.com/ncbi/sra-tools). The adapter sequences were removed (if not removed already) from the datasets using fastp. Additionally, 3 nucleotides from the 5' end of all reads were also trimmed using fastp as these nucleotides were generally found to be of a low-quality score. Reads that aligned to non-coding RNA sequences (rRNA, tRNA, snRNA, snoRNA, and miRNA) were removed using Bowtie2 (version: 2.3.4.1). The FASTQ files were then aligned with a list of protein-coding mRNAs to create BAM (Binary alignment map) files.

### Mapping of ribosome profiling (ribo-seq) reads onto the coding sequence (CDS), the ISR and the 3′UTR of mRNAs

We first analyzed the length distribution of the ribo-seq reads in each dataset. Based on this distribution, we chose the reads of 3 most abundant lengths for further analyses. Ribo-seq reads were assigned to different regions of an mRNA (i.e., CDS, ISR and 3′UTR) based on the alignment of the beginning of the read to any of these regions (Fig S1). To avoid ambiguity during the assignment of the ribo-seq reads to different regions of an mRNA, we followed these criteria:

i. For coding sequence (CDS): reads that align to the region from 12 nucleotides upstream of the start codon till 22^nd^ nucleotide upstream of the canonical stop codon.
ii. For ISR: reads that align to the region from 12 nucleotides upstream of the canonical stop codon till 22^nd^ nucleotide upstream of the downstream in-frame stop codon.
iii. For 3′UTR: reads that align to the region from 12 nucleotides upstream of the downstream in-frame stop codon till the end of the mRNA. Only those reads that showed 100% alignment to an mRNA region were considered (Even a single mismatch was not allowed).

### Identification of potential SCR candidates

#### Selection based on ribo-seq read density

The mRNAs with higher density of reads in their ISR than that in the coding sequence were removed from the analysis as this feature is not consistent with SCR. On the contrary, mRNAs with a 4-fold higher density of reads in their ISR than that in the rest of the 3′UTR were included as this is a signature of translational readthrough.

#### Selection based on ribo-seq read coverage

To increase the stringency of this screening process, we applied three more selection criteria based on the coverage:

i. at least 30 ribo-seq reads should map onto the ISR of a given mRNA
ii. > 50% of ISR and < 25% 3′UTR should be covered by ribo-seq reads
iii. there should be at least one ribo-seq read spanning the canonical stop codon.

#### Selection based on three-nucleotide periodicity

We looked at a 62-nucleotide window length around the start and the stop codons to ensure that the ribo-seq datasets showed three-nucleotide periodicity. To quantify these frame biases, all genes with at least 200 reads in the coding sequence were considered. Reads were assigned to three frames based on which frame the first nucleotide aligns with on an mRNA sequence. We computed the mean and the standard deviation of the fraction of reads that fell in each frame across all the codons of the CDS region. This was then used as the reference distribution against which each of the SCR candidates (filtered based on density and coverage) was compared. The candidates which satisfy the following criterion were selected: the fraction of reads that fell in each frame in the CDS and the ISR regions (test distributions) are within two standard deviations of the reference distribution, in at least one frame.

All codes used in this study are available at: https://github.com/Divyoj-Singh/Stop_codon_readthrough_pipeline

### Experimental validation

#### Plasmid constructs

Luciferase constructs for luminescence-based SCR assay were generated in pcDNA 3.1 backbone. The coding sequence of the test gene along with the canonical stop codon and the ISR was cloned upstream of and in‐frame with the coding sequence of the firefly luciferase (FLuc) between *Hind*III and *BamH*I sites (*MUB6* and *RPS15AD*) or *Kpn*I and *BamH*I sites (*CAM1* and *CURT1B*). A linker sequence (GGCGGCTCCGGCGGCTCCCTCGTGCTCGGG) was included upstream of the FLuc coding sequence.

#### In vitro transcription and translation

The plasmid DNA was linearized using *Not*I enzyme, and 2 µg of the linearized DNA was transcribed *in vitro* using T7 RNA polymerase (Thermo Fisher Scientific). The resultant RNA was purified using GeneJET RNA purification kit (Thermo Fisher Scientific). The concentration and quality of the RNA were measured using BioPhotometer (Eppendorf). 2-3 μg of the purified RNA was *in vitro* translated using wheat germ extract (Promega) at 25 °C for 2 h as per the manufacturer’s instructions. Luciferase activity was then measured using the Luciferase Assay System (Promega Corporation) in the GloMax Explorer System (Promega Corporation).

## Acknowledgements

We thank Prof. Utpal Nath, Indian Institute of Science, for providing *A. thaliana* tissue samples. We acknowledge the financial support from the Department of Biotechnology (DBT) – Wellcome Trust India Alliance Intermediate Fellowship (IA/I/15/1/501833), Swarnajayanti Fellowship (SB/SJF/2020-21/18) from the Department of Science and Technology (DST), STARS grant from the Ministry of Education, DST Funds for Improvement of S&T infrastructure, DBT - Indian Institute of Science Partnership Program for Advanced Research in Biological Sciences, and funds from the University Grants Commission (UGC), India. S.S. and D.S. are supported by KVPY fellowship awarded by DST.

## Author contributions

**SS:** Conceptualization, Data Curation, Formal Analysis, Investigation, Methodology, Visualization, Software. **DS:** Data Curation, Investigation, Methodology, Visualization, Software. **AS:** Investigation, Methodology. **SME**: Conceptualization, Funding Acquisition, Formal analysis, Visualization, Project Administration, Resources, Supervision, Writing – Original Draft Preparation.

## Legends to supplementary figures

**Figure S1.**
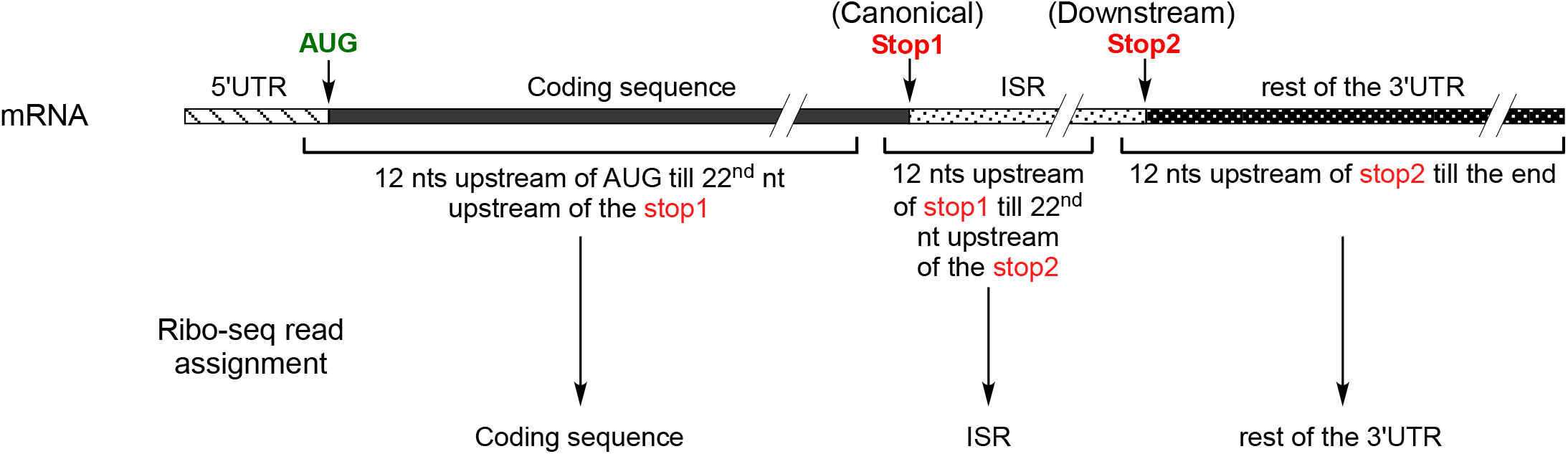
Schematic to explain the assignment of ribo-seq reads to various regions of an mRNA.

**Figure S2.**
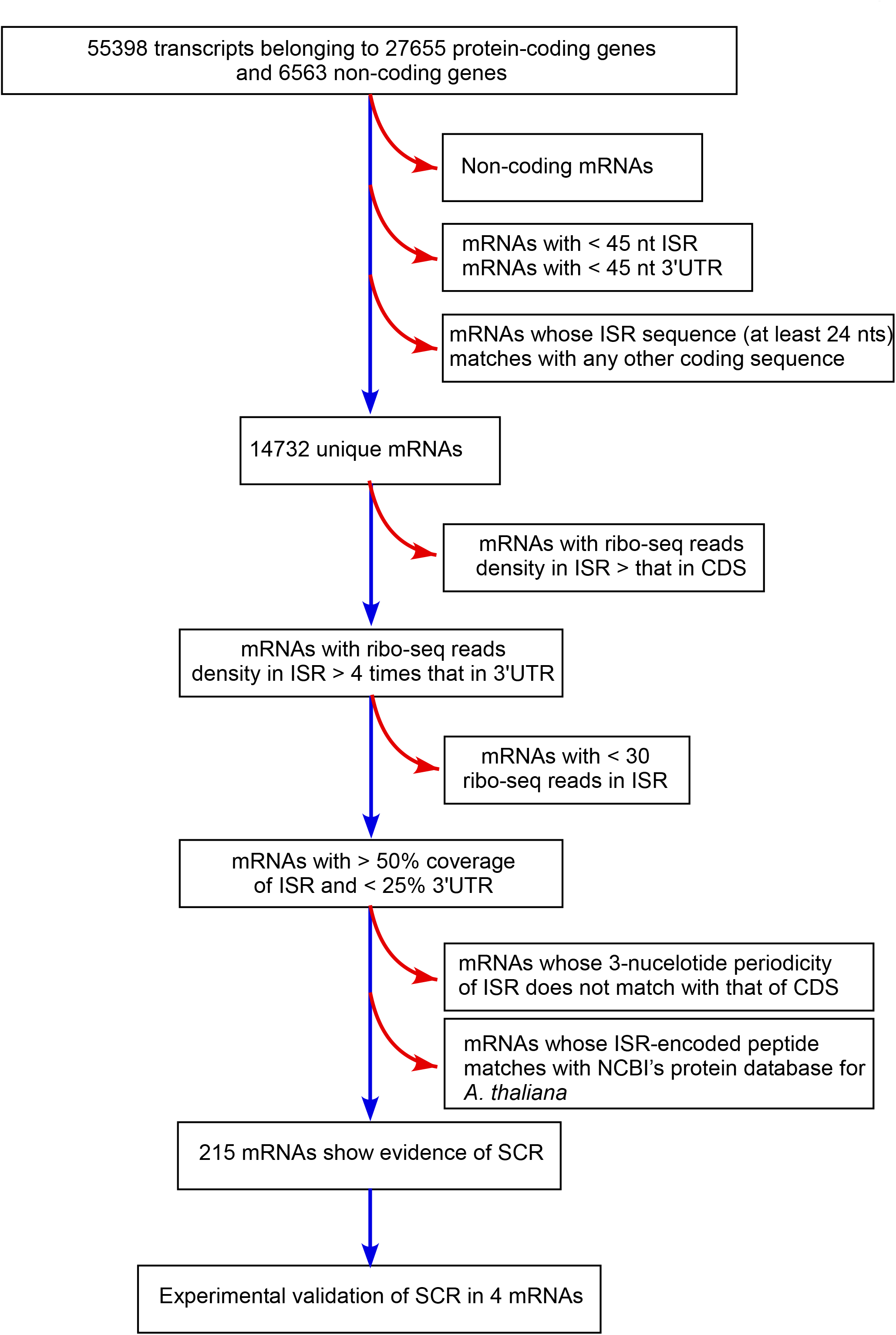
Flow chart showing the four-level screening method to identify mRNAs that show SCR in *A. thaliana.* Blue arrows indicate inclusion and red arrows indicate exclusion.

